# IL-38 regulates intestinal stem cell homeostasis by inducing WNT signaling and beneficial IL-1β secretion

**DOI:** 10.1101/2023.04.04.535251

**Authors:** Alberto Dinarello, Makenna May, Jesus Amo-Aparicio, Tania Azam, Joseph M Gaballa, Carlo Marchetti, Annachiara Tesoriere, Rachele Ghirardo, Jasmina S Redzic, William Webber, Shaikh M Atif, Suzhao Li, Elan Z Eisenmesser, Dennis M de Graaf, Charles A Dinarello

**Affiliations:** Department of Medicine, University of Colorado Anschutz Medical Campus, Aurora, CO 80045, USA; Department of Biology, University of Padova, Padova, 35121, Italy; Department of Biochemistry and Molecular Genetics, School of Medicine, University of Colorado Denver, 12801 E 17th Ave, Aurora, CO, 80045, USA; Institute of Innate Immunity, University Hospital Bonn Biomedical Center, 53127 Bonn, Germany

## Abstract

The IL-1 Family member IL-38 has been characterized primarily as an anti-inflammatory cytokine in human and mouse models of systemic diseases. Here, we examined the role of IL-38 in the murine small intestine (SI). Immunostaining of SI revealed that IL-38 expression partially confines to intestinal stem cells. Cultures of intestinal organoids reveal IL-38 functions as a growth factor by increasing organoid size via inducing WNT3a. In contrast, organoids from IL-38 deficient mice develop more slowly. This reduction in size is likely due to downregulation of intestinal stemness markers (i.e., *Fzd5*, *Ephb2*, *Olfm4*) expression compared with wild type organoids. IL-38 binding to IL-1R6 is postulated to recruit the co-receptor IL-1R9. Therefore, to analyze the molecular mechanisms of IL-38 signaling, we also examined organoids from IL-1R9 deficient mice. Unexpectedly, these organoids, although significantly smaller than wild type, respond to IL-38, suggesting that IL-1R9 is not involved in IL-38 signaling in the stem cell crypt. Nevertheless, silencing of IL-1R6 disabled the organoid response to the growth property of IL-38, thus suggesting IL-1R6 as the main receptor used by IL-38 in the crypt compartment. In organoids from wild type mice, IL-38 stimulation induced low concentrations of IL-1β which contribute to organoid growth. However, high concentrations of IL-1β have detrimental effects on the cultures that were prevented by treatment with recombinant IL-38. Overall, our data demonstrate an important regulatory function of IL-38 as a growth factor, and as an anti-inflammatory molecule in the SI, maintaining homeostasis.

**Significance:** The IL-1 family member IL-38 has been characterized primarily as an anti-inflammatory cytokine for systemic diseases. Here we describe a fundamental role of IL-38 in driving intestinal stem cell differentiation through the upregulation of WNT3a and IL-1β. Our findings reveal a dual role of IL-38 in regulating intestinal functions; (a) in resting conditions IL-38 maintains intestinal homeostasis, driving WNT3a production and organoid budding, whereas (b) in highly inflamed conditions, IL-38 contributes to proper recovery, by exerting anti-inflammatory activities. Thus, we demonstrate a pivotal role of IL-38 in driving tissue turnover and maintenance of homeostasis in intestinal health.

## Introduction

IL-38 is a cytokine that belongs to the IL-1 family (1). It is encoded by *Il1f10* gene and is expressed in infiltrating immune cells of colonic samples from patients with inflammatory bowel disease (IBD), particularly CD19^+^ B cells (2) and CD123^+^ cells (3). IL-38 is mostly studied as an anti-inflammatory cytokine (1, 4) and exerts protective functions in several organs, suggesting that this protein mainly works in the maintenance of organ homeostasis (1). As demonstrated by Van de Veerdonk and collaborators, IL-38 has biological effects in immune cells that resembles IL-36Ra functions, downregulating the levels of IL-22, IL-17, and IFNγ in *Candida*-stimulated Th17 cells (5). A recent study by The and collaborators also demonstrated that IL-38 dampens aortic valve calcification, probably dependent on IL-1R9 (encoded by *Il1rapl1* gene), inhibiting NLR family pyrin domain containing 3 (NLRP3) and IL-1β maturation (6). Moreover, *Il1f10* deficient mice exhibit higher levels of inflammation when exposed to DSS treatment compared to wild type mice and express higher levels of NLRP3 and caspase-1 (7). Regarding IBD, IL-38 is expressed in colon tissue of ulcerative colitis patients compared to healthy controls (8), reduces inflammation in the colon (2).

Although most studies demonstrate that IL-38 has anti-inflammatory properties by, for example, blocking IL-1β maturation, the biological functions of IL-38 are more complex than only inhibiting the inflammatory response and some *in vivo* studies demonstrate that the lack of this cytokine favors diseases progression. For instance, as shown in Huard *et al*. (9), IL-38 ablation in mice ameliorates autoimmune encephalomyelitis leading to downregulation of inflammation markers like *Mertk*, *Tnfa*, *Ptgs2*, *Tgfb1* and *Tgfb2* (9). On the other hand, experiments performed on mouse and A431 human epidermoids cancer cell line reveal that IL-38 has a pro-tumorigenic function and can stimulate cancer cell proliferation using an IL-1R6-dependent mechanism (10). Moreover, as regards keratinocytes differentiation, Mermoud and collaborators (11) have recently demonstrated that IL-38 is expressed in primary keratinocytes as well as in the N/TERT1 keratinocyte cell line. Notably, the overexpression of IL-38 promotes the differentiation of these cells, inhibiting proliferation (11).

Another debated aspect of IL-38 biology consists in its receptor. As previously proposed by van de Veerdonk and collaborators (5), IL-38 binds IL-36R (IL-1R6), encoded by *Il1rl2* gene. Other pieces of evidence regarding the interaction of IL-38 with IL-1R6 come from Mora and collaborator’s experiments (12), who demonstrated that IL-38 binds either IL-1R6 and IL-1R9 (12). Additionally, pulldown experiments performed by Zhou and coworkers (10) support the IL-38-IL-1R6 interaction. IL-38 may also bind IL-1R1 and the affinity of interaction depends on the truncation of the N-terminal amino acids of IL-38 (13).

Given the emerging importance of a therapeutic role for IL-38 (14, 15, 16) and the function of this cytokine in the intestine (2, 7, 8, 17), here we analyzed the effects of IL-38 in intestinal stem cells (ISCs) in organoid cultures derived from wild type and *Il1f10* mice. Moreover, to better dissect the mechanism of action of IL-38 in the gut, we employed *Il1rapl1* deficient mice and observed that IL-1R9 is not essential for a response to IL-38. Additionally, silencing experiments highlight the role of IL-1R6 in IL-38 signaling. Moreover, we discovered that IL-38 induces low concentrations of IL-1β. This discovery, confirmed in *Il1r1* and *Nlrp3* deficient mice, reveals that homeostatic levels of IL-1β are beneficial for organoids.

## Results

### 1. *Il1f10* deficient intestine show lower expression of intestinal stem cell markers

Located in the intestinal crypts, ISCs represent the precursor cells of the entire intestine (18). These cells regulate homeostasis of the intestine, producing soluble factors that affect the fate of the intestine. The crypts are divided into two zones: the stem cell zone and the transient applying zone (TA). The stem cell zone contains the crypt base columnar cells that act as ISCs, as well as Paneth cells that protect and feed the ISCs (19). The TA zone is composed by lineage-committed stem cells, which rapidly divide and differentiate; the mature cell zone is populated by large numbers of epithelial cells (19). To determine these areas, cell markers have been identified to show differentiated and undifferentiated cells. Historically, the most important marker of ISCs is LGR5 (18, 20, 21, 22), a G-protein Coupled Receptor and a target of the WNT pathway (23). As demonstrated by Carmon and collaborators, LGR5 binds R-spondins for enhancing the activation of the WNT/β-catenin signaling in WNT responsive cells (24). The WNT/β-catenin axis is fundamental for crypt homeostasis and maintains ISCs in an undifferentiated state (21); however, WNT/β-catenin signaling can also drive the differentiation of ISCs in Paneth cells (25).

IL-38 is an anti-inflammatory cytokine that can protect the organism challenged with highly inflammatory triggers, as recently observed in dextran sodium sulfate-colitis or aortic calcification models (6,7). We sought to confirm whether IL-38 is expressed in the mouse intestine (8) and if the IL-38 protein is expressed in the small intestine at the level of the stem cell crypts. As shown in Figure 1, IL-38 staining co-localizes with LGR5 staining, revealing for the first time that IL-38 is expressed in stem cell crypts and suggesting a role of this cytokine in intestinal stem cells.

**Figure 1:**
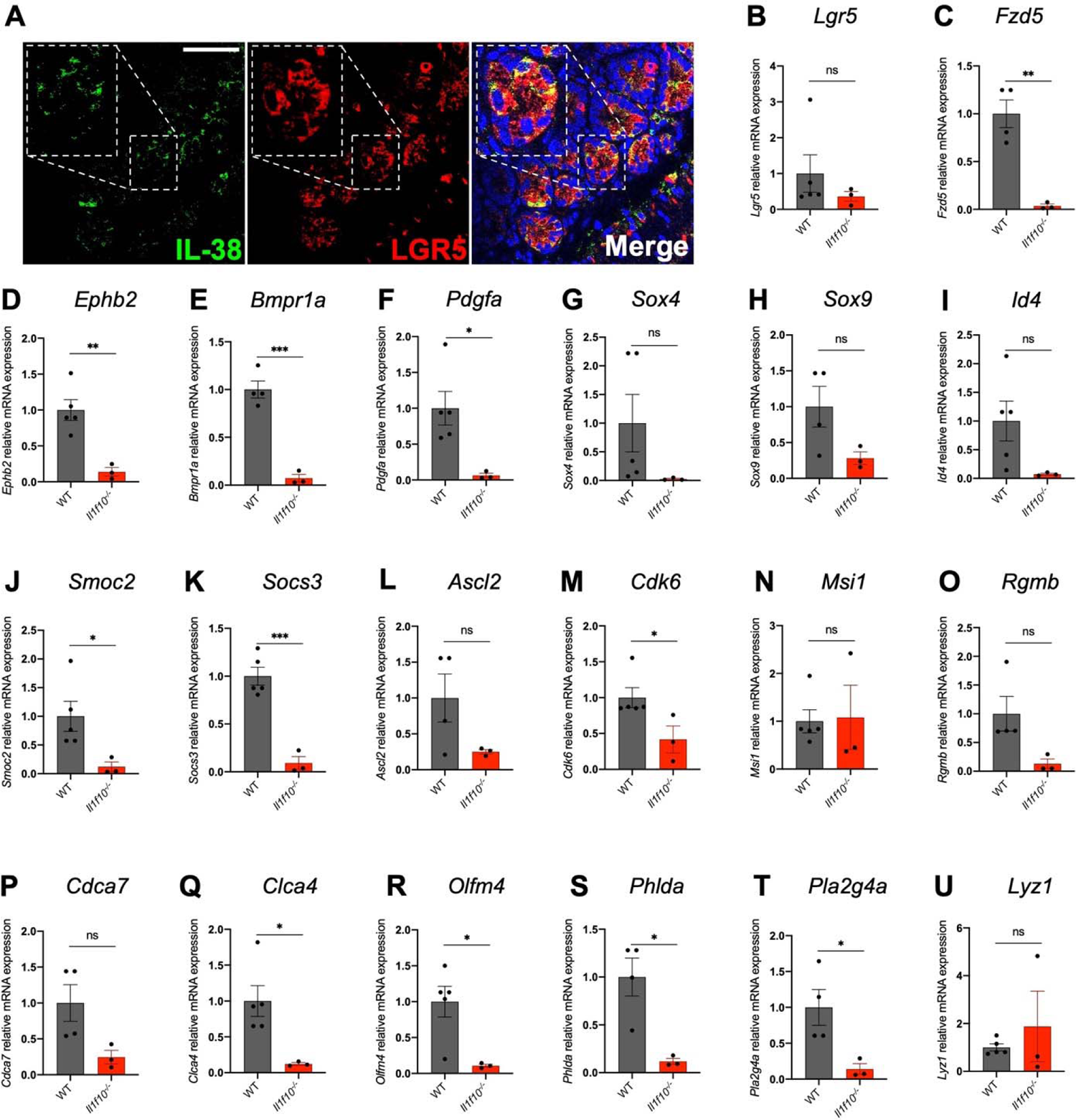
IL-38 is essential for intestinal crypt gene expression. (**A**) Staining of IL-38 (green) and LGR5 (red) in wild type intestine. An enlargement of one intestinal crypt is shown in dashed frame. Scale bar = 50 μm. (**B-U**) gene expression analysis of *Lgr5* (**B**), *Fzd5* (**C**), *Ephb2* (**D**), *Bmpr1a* (**E**), *Pdgfa* (**F**), *Sox4* (**G**), *Sox9* (**H**), *Id4* (**I**), *Smoc2* (**J**), *Socs3* (**K**), *Ascl2* (**L**), *Cdk6* (**M**), *Msi1* (**N**), *Rgmb* (**O**), *Cdca7* (**P**), *Clca4* (**Q**), *Olfm4* (**R**), *Phlda* (**S**), *Pla2g4a* (**T**) and *Lyz1* (**U**) in intestines of wild type and *Il1f10* deficient mice. *p < 0.05; **p < 0.01, ***p < 0.001, ns = not significant. Mean ± SEM.

To confirm the putative contribution of IL-38 to homeostasis of intestinal stem cells, we measured the level of expression of stem cell niche markers (taken from (26) and (27)) in intestines from wild type and *Il1f10* deficient mice. Among the markers analyzed, *Fzd5, Ephb2, Bmpr1a, Clca4, Pdgfa, Olfm4, Smoc2, Socs3, Cdk6, Phlda* and *Pla2g4a* are significantly downregulated in *Il1f10* deficient mice compared to wild type (Figure 1B-U), demonstrating the pivotal role of IL-38 in intestinal stem cell niche gene expression. These findings are further corroborated by western blot of intestinal extracts against LGR5. As shown in Figure S1, LGR5 expression is significantly reduced in *Il1f10* deficient intestines compared to wild type.

### 2. IL-38 is a growth factor in intestinal organoids

To test the functions of IL-38 *ex vivo*, we generated organoid cultures from wild type SI. To visualize the presence of endogenous IL-38 in these organoids, we performed immunofluorescence staining for IL-38 (Green) and Lyz1 (Red) to mark Paneth cells. As observed in Figure 2A, IL-38 seems to be ubiquitously expressed in organoids. Organoids were then stimulated with recombinant mouse IL-38 at three different concentrations. Interestingly, 2 ng/ml and 200 ng/ml did not affect organoid size (Figure S1B-E), whereas 20 ng/ml of IL-38 increased the size of organoids (Figure 2B,C). Additionally, we injected recombinant mouse IL-38 in wild type mice every day for 15 days and collected intestines to generate organoid cultures. Notably, organoids obtained from IL-38 injected mice are larger than vehicle injected mice, confirming that IL-38 can function as a growth factor in healthy *in vivo* intestines (Figure 2D,E).

**Figure 2:**
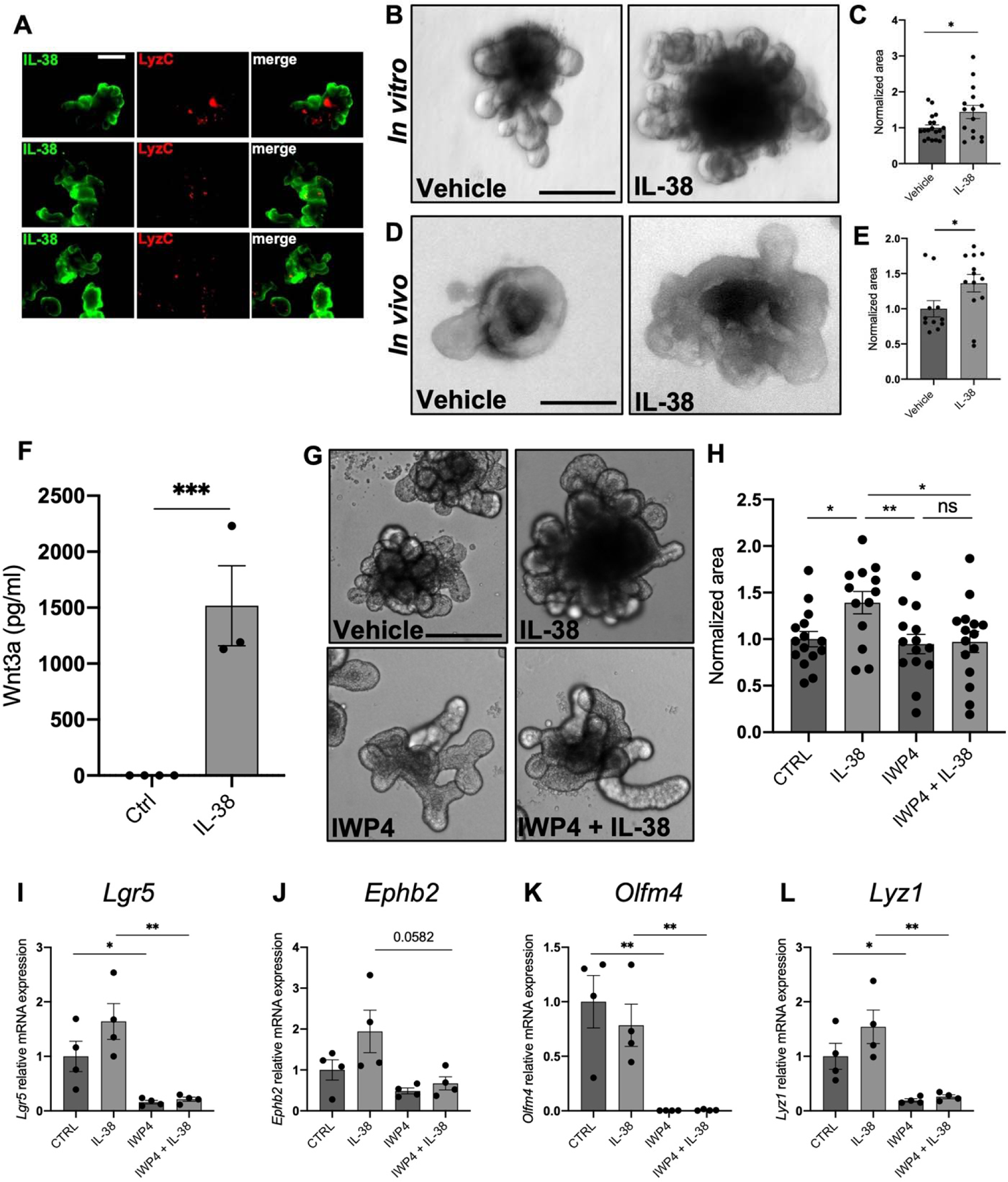
IL-38 works as a growth factor stimulating WNT3a secretion. (**A**) Ubiquitous expression of IL-38 in intestinal organoids: immunofluorescence staining for IL-38 (green) and LyzC (red) to mark Paneth cells. (**B**) Representative pictures and (**C**) measurement of wild type organoids treated with 20 ng/ml IL-38 for 6 days. Scale bar: 200 μm. (**D**) Representative pictures and (**E**) measurements of wild type organoids obtained from vehicle- and IL-38-injected mice. Scale bar: 200 μm. (**F**) WNT3a expression in the supernatants in wild type organoids treated with 20 ng/ml IL-38 for 6 days. (**G**) Representative pictures and (**H**) measurement of wild type organoids treated with vehicle, 20 ng/ml IL-38, 5 μM IWP4, and 5 μM IWP4 + 20 ng/ml IL-38. Scale bar: 200 μm. (**I-L**) Gene expression of *Lgr5* (**I**)*, Ephb2* (**J**)*, Olfm4* (**K**), and *Lyz1* (**L**) in wild type organoids treated with vehicle, 20 ng/ml IL-38, 5 μM IWP4, and 5 μM IWP4 + 20 ng/ml IL-38. *p < 0.05; **p < 0.01, ***p < 0.001, ns = not significant. Mean ± SEM.

WNT3a is the major soluble factor produced by Paneth cells that drive ISCs differentiation (24). To understand how IL-38 affects organoid growth, we measured the levels of WNT3a in organoid. As shown in Figure 2F, we observed a significant increase with IL-38 stimulation (1500-fold increase, p < 0.001) (Figure 2F). To confirm whether the increase of organoid size depends on IL-38-induced WNT3a secretion, we treated organoids with the WNT inhibitor IWP4 (28) either with or without IL-38 stimulation. As expected, IL-38 stimulated organoid growth, whereas IWP4 did not affect organoid size compared to vehicle yet changed organoid shape (Figure 2G,H). Moreover, organoids treated with IL-38 and IWP4 combination do not show significant differences when compared to either vehicle or IWP4-treated organoids but are significantly smaller compared to IL-38-treated organoids (Figure 2G,H). Furthermore, we observed the same phenotype in organoids treated with IWP4 and IL-38+IWP4 (Figure 2G,H). These data suggest that IL-38 stimulates organoid growth by inducing WNT3a production, and that these effects are dependent on the WNT pathway. To confirm the actual inhibition of WNT pathway, we also measured the expression of WNT and stemness marker genes and we observed that *Lgr5*, *Lyz1* and *Olfm4* are significantly downregulated in IWP4-treated organoids compared to control and IL-38-treated organoids (Figure 2I-L).

### 3. *Il1f10* deficient (*Il1f10^-/-^*) organoids show growth defects

Organoid cultures from wild type and IL-38 deficient mice were evaluated for growth. IL-38 deficient organoids are significantly smaller than wild type organoids and, as expected, 20 ng/ml recombinant IL-38 significantly increased the size of either wild type or IL-38 deficient organoids (Figure 3A,B).

**Figure 3:**
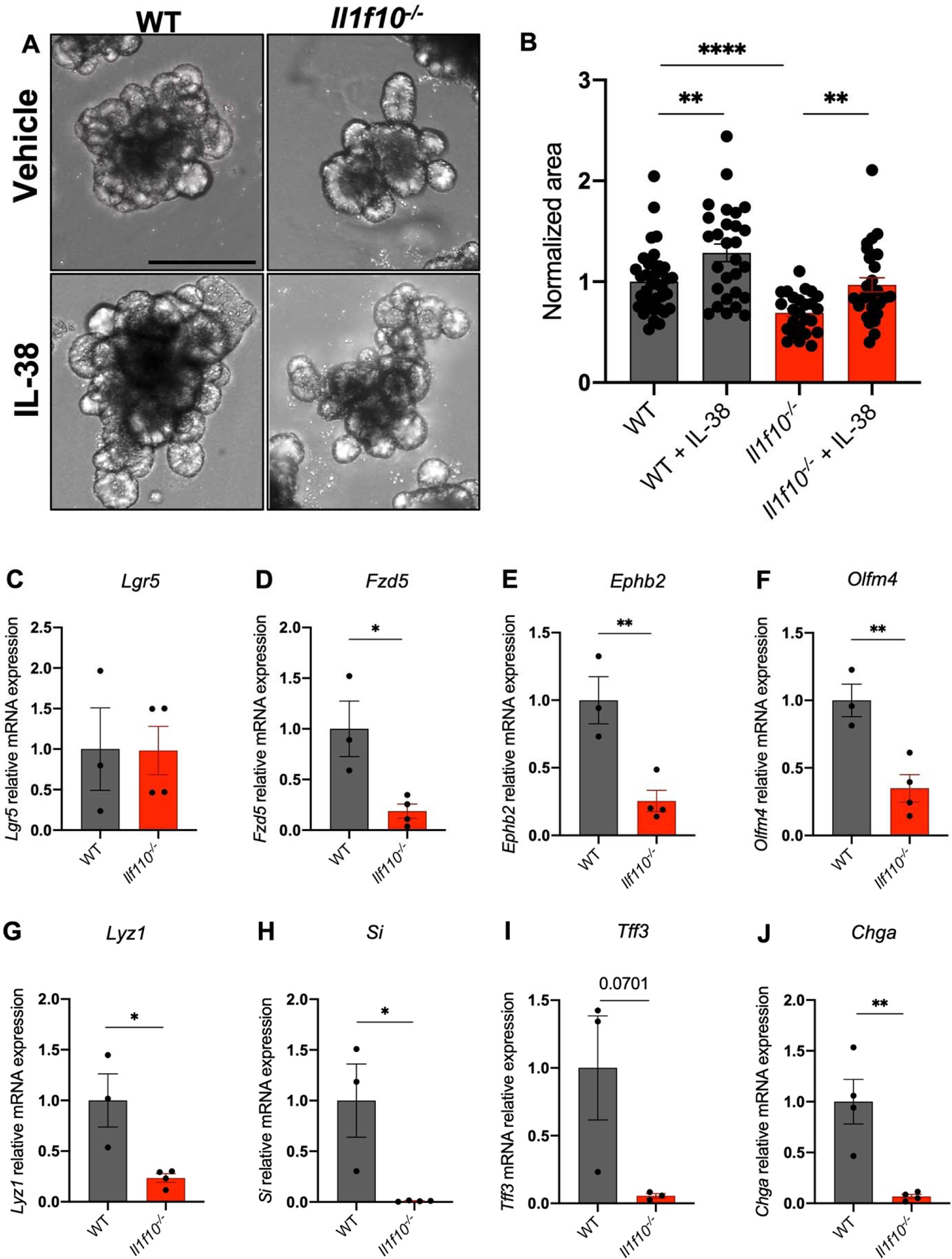
*Il1f10* deficient organoids show growth defects. (**A**) Representative pictures and (**B**) measurements of wild type and *Il1f10^-/-^* organoids treated either with vehicle or 20 ng/ml IL-38 for 6 days. Scale bar: 200 μm. (**C-J**) Gene expression analysis of *Lgr5* (**C**)*, Fzd5* (**D**)*, Ephb2* (**E**)*, Olfm4* (**F**), *Lyz1* (**G**), *Si* (**H**), *Tff3* (**I**) and *Chga* (**J**) in wild type and *Il1f10^-/-^*organoids. *p < 0.05; **p < 0.01, ****p < 0.0001. Mean ± SEM.

Interestingly, we observed a marked downregulation of stemness markers in *Il1f10* knockout organoids compared to wild type. *Lgr5* does not show significant differences between *Il1f10^-/-^* and wild type organoids, whereas *Fzd5*, *Ephb2* and *Olfm4* are significantly downregulated in *Il1f10* deficient organoids when compared to wild type (Figure 3C-F)

To further characterize *Il1f10^-/-^* organoids, we measured the levels of expression of markers of other cell types such as *Lyz1* (Paneth cells), *Si* (absorptive cells), *Tff3* (goblet cells) and *Chga* (goblet cells) (29). Notably, all these markers were downregulated in *Il1f10* deficient organoids, suggesting that the lack of IL-38 fundamentally affects the differentiation of ISC through several intestinal cell types (Figure 3G-J).

### 4. *Il1rapl1* deficient (*Il1rapl1^-/y^*) organoids show growth defects

IL-38 functions are determined by its interaction with its putative receptors IL-1R6 and IL-1R9 (5, 10, 12, 13).

Using RNA extracted from wild type and *Il1rapl1* deficient intestines, we demonstrated that, among all the transcripts analyzed in Figure 1, *Fzd5, Bmpr1a, Cdca7* and *Phlda* are significantly downregulated in *Il1rapl1* deficient intestines compared to wild type controls (Figure 4A-D). In contrast, other genes are not differentially expressed between wild type and *Il1rapl1* deficient intestines, whereas *Clca4* resulted significantly upregulated (Figure S2A-P). These observations suggest that IL-1R9 is involved in the correct intestinal stem cell niche homeostasis, but its absence impacts stemness markers less than IL-38 deficiency.

**Figure 4:**
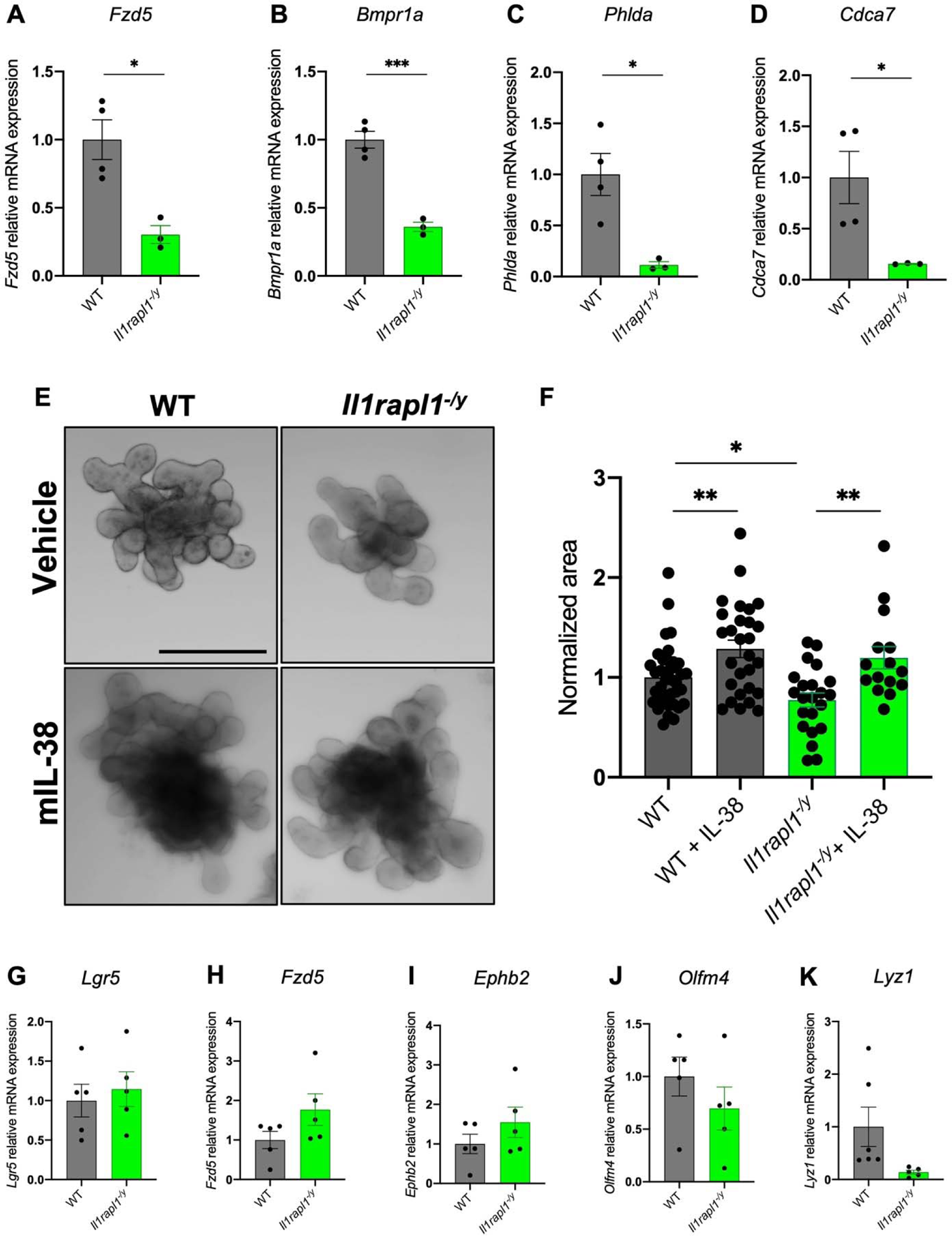
*Il1rapl1* deficient organoids show growth defects. (**A-D**) Gene expression analysis of *Fzd5* (**A**)*, Bmpr1a* (**B**)*, Phlda* (**C**) and *Cdca7* (**D**) in intestines of wild type and *Il1rapl1* deficient mice. (**E**) Representative pictures and (**F**) measurements of wild type and *Il1rapl1* deficient organoids treated either with vehicle or 20 ng/ml IL-38 for 6 days. Scale bar: 200 μm. (**G-K**) Gene expression analysis of *Lgr5* (**G**)*, Fzd5* (**H**)*, Ephb2* (**I**)*, Olfm4* (**J**) and *Lyz1* (**K**) in wild type and *Il1rapl1* deficient organoids. *p < 0.05; **p < 0.01. Mean ± SEM.

Next, we cultured organoids derived from wild type and *Il1rapl1* deficient mice and we observed that *Il1rapl1* knockout organoids are significantly smaller than wild type-derived organoids, highlighting the importance of IL-1R9 in intestinal stem cell niche homeostasis (Figure 4E,F). Moreover, IL-38 stimulation of organoid cultures revealed that both wild type and *Il1rapl1* deficient cultures respond to stimulation. In fact, we observed a significant increase of organoid size in IL-38-treated wild type and *Il1rapl1* deficient organoids compared to untreated controls (Figure 4E,F). In addition, RT-qPCR for *Lgr5, Fzd5, Ephb2, Olfm4* and *Lyz1* revealed no differences between wild type and *Il1rapl1* KO organoids (Figure 4G-K). These data demonstrate that IL-1R9 is not required for IL-38 biological activities in intestinal organoids and that another receptor is involved in IL-38-signaling.

### 5. IL-1R6 is involved in IL-38 signaling in the intestinal crypt

IL-1R6 is encoded by the *Il1rl2* gene. In order to understand whether the lack of this receptor has a detrimental effect on organoid growth, we used *Il1rl2* siRNAs on wild type organoids. As shown in Figures 5 A,B, *Il1rl2* silencing determines a significant reduction of organoid size, suggesting an important role of IL-1R6 in this process. After validating *Il1rl2* RNA silencing by RT-qPCR (Figure 5C), we measured the expression levels of *Lgr5, Fzd5, Ephb2, Olfm4* and *Lyz1.* Differently to what we observed with *Il1rapl1* deficient organoids, *Il1rl2* silencing resulted in a significant downregulation of *Fzd5* and *Ephb2*, without significantly affecting the expression of the other markers analyzed (Figure 5D-H).

**Figure 5:**
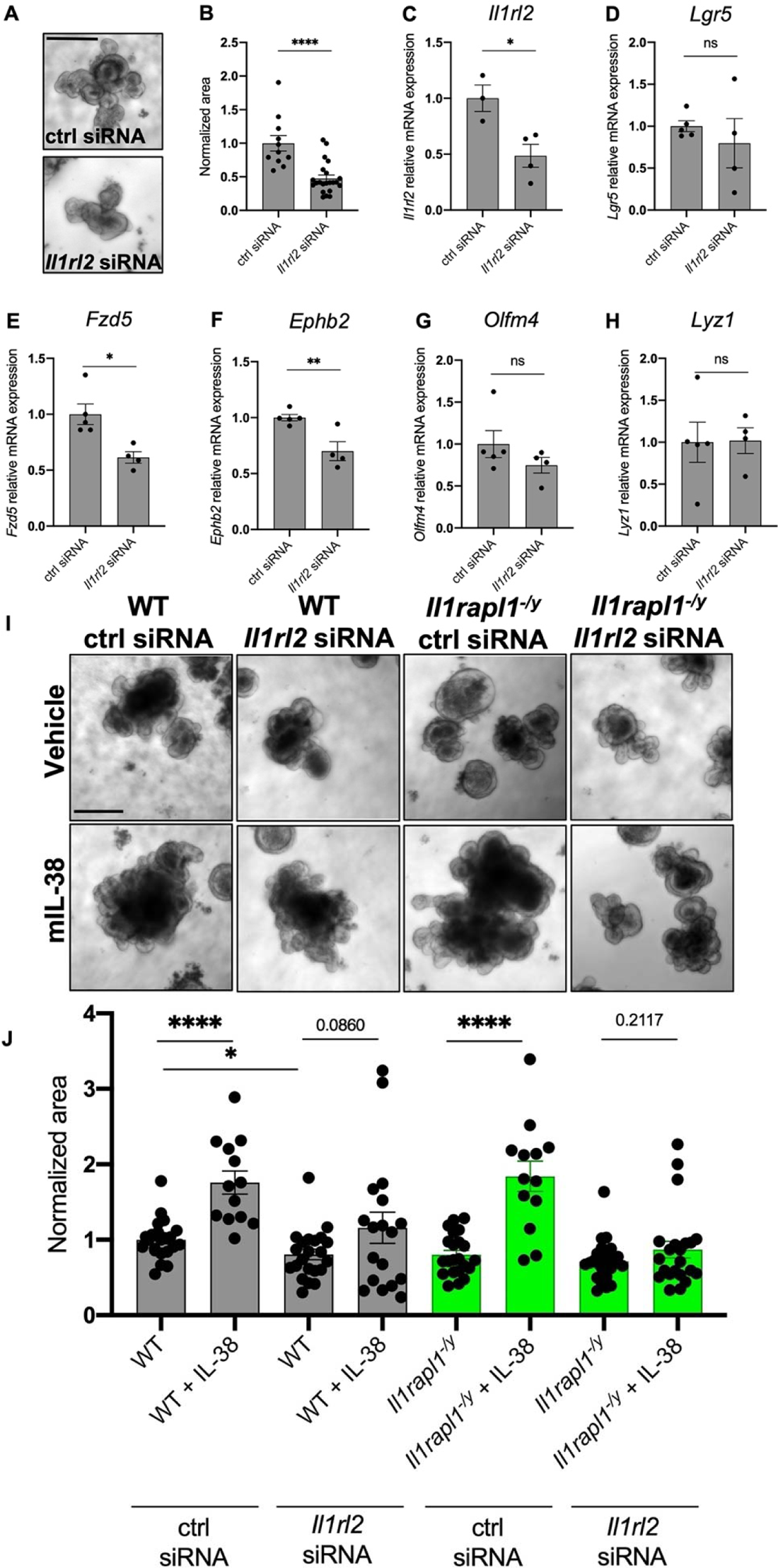
*Il1rl2* silencing negatively affects organoid growth. (**A**) Representative pictures and (**B**) measurements of wild type organoids treated with control siRNA or *Il1rl2* siRNA for 6 days. (**C-H**) Gene expression analysis of *Il1rl2* (**C**), *Lgr5* (**D**)*, Fzd5* (**E**)*, Ephb2* (**F**)*, Olfm4* (**G**) and *Lyz1* (**H**) in wild type organoids treated with either control siRNA or *Il1rl2* siRNA for 6 days. (**I**) Representative pictures and (**J**) measurements of wild type and *Il1rapl1* deficient organoids treated with control siRNA, *Il1rl2* siRNA, vehicle or IL-38 for 6 days. Scale bar: 200 μm. *p < 0.05; **p < 0.01; ****p < 0.0001. Mean ± SEM.

Next, we sought to assess whether organoids can respond to IL-38 even in the absence of IL-1R6. To do so, we incubated wild type organoids with either control siRNA or *Il1rl2* siRNA and we stimulated them for 6 days with IL-38. As observed in Figure 5I and J, recombinant IL-38 significantly increases the organoids size in control siRNA treated organoids (p < 0.0001), but we could not see significant differences between *Il1rl2* silenced organoids treated with vehicle or with recombinant IL-38 (p = 0.086), suggesting that IL-38 increases organoids growth through IL-1R6. To better evaluate the role of IL-1R6 and IL-1R9 in IL-38 biological activity, we silenced *Il1rl2* in *Il1rapl1* deficient organoids. As previously observed in Figure 4E and F, *Il1rapl1* deficient organoids indeed respond to IL-38, however, upon silencing with *Il1rl2* siRNA, IL-38 did not stimulate growth (Figure 5I,J). These data demonstrate that IL-38 signaling depends on IL-1R6 whereas IL-1R9 plays a separate role in intestinal organoid homeostasis.

### 6. Low concentrations of IL-1β stimulate organoid growth

Together with the induction of WNT3a in organoid supernatants (as observed in Figure 2), IL-38 induces a low but significant concentration of IL-1β (about 25 pg/ml) (Figure 6A), suggesting a possible role of this cytokine in the correct homeostasis of organoids. To test the importance of IL-1β in intestinal organoid cultures we first decided to block IL-1 signaling by treating wild type organoids with 10 μg/ml Anakinra (a commercially available antagonist of IL-1 receptor) for 6 days. Anakinra significantly reduces organoids size (p < 0.0001), suggesting the importance of IL-1 signaling for the proper development of intestinal cells (Figure 6B,C). To confirm this result, we generated cultures of *Il1r1* knockout organoids. As observed in Figure 6D,E, *Il1r1* deficient organoids are significantly smaller than wild type (p < 0.0001), recapitulating with a genetic model what we previously observed with Anakinra treatment (Figure 6B-E). Similarly, *Nlrp3* knockout organoids, that do not produce NLRP3, fundamental for inflammasome formation thus for the conversion of pro-IL-1β into mature IL-1β (30), show a marked reduction in size when compared to wild type (p < 0.0001) (Figure 6F,G). These data demonstrate a central role of IL-1β in organoid growth.

**Figure 6:**
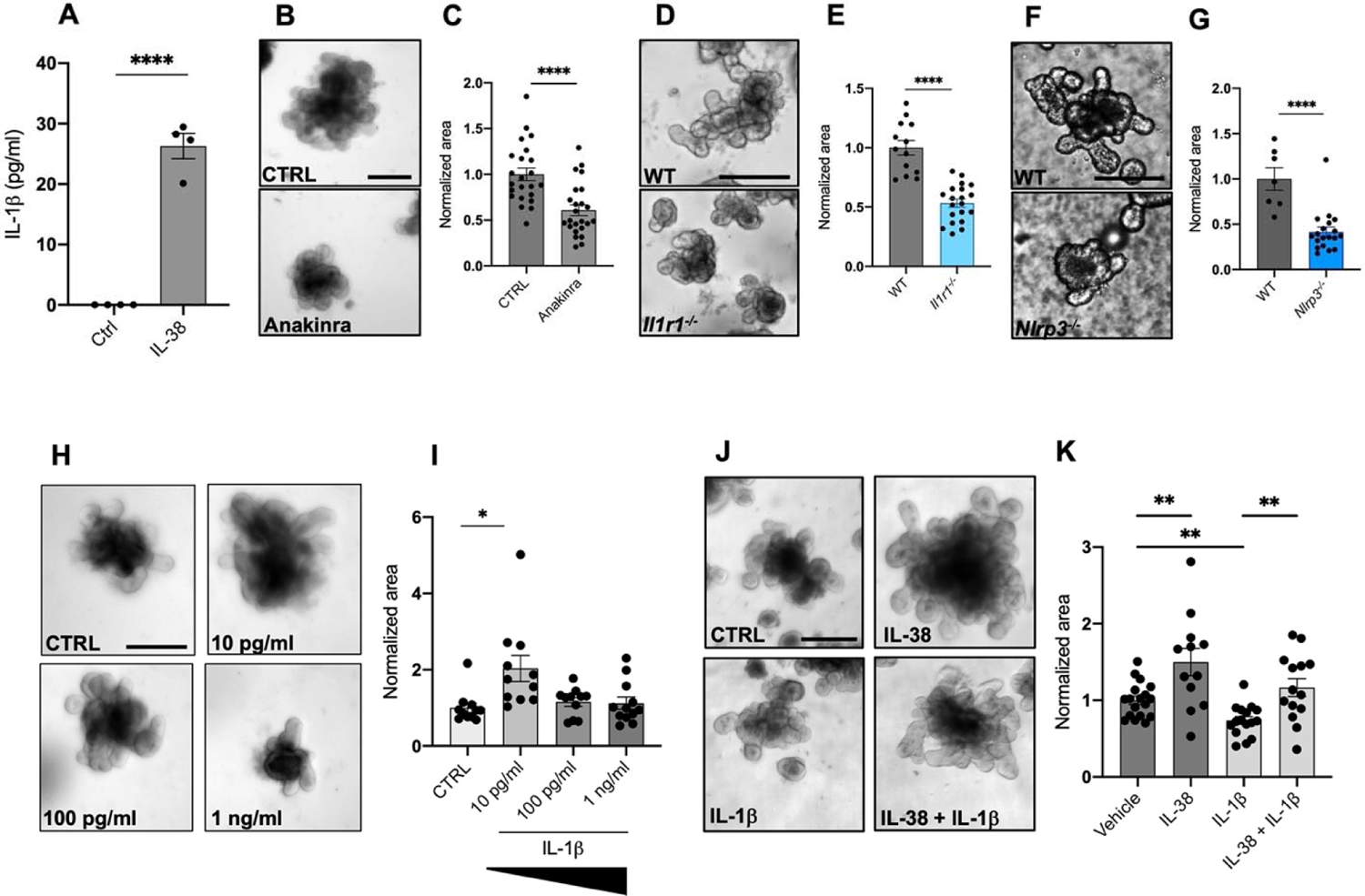
Low concentrations of IL-1β induce organoid growth. (**A**) IL-1β expression in the supernatants in wild type organoids treated with 20 ng/ml IL-38 for 6 days. (**B**) Representative pictures and (**C**) measurements of wild type organoids treated with 10 μg/ml Anakinra for 6 days. Scale bar: 200 μm. (**D**) Representative pictures and (**E**) measurement of wild type and *Il1r1^-/-^* organoids. Scale bar: 200 μm. (**F**) Representative pictures and (G) measurement of wild type and *Nlrp3^-/-^* organoids. Scale bar: 200 μm. (**H**) Representative pictures and (**I**) measurements of wild type organoids treated with 10 pg/ml, 100 pg/ml and 1 ng/ml IL-1β for 6 days. Scale bar: 200 μm. (**J**) Representative pictures and (**K**) measurements of wild type organoids treated with vehicle, 20 ng/ml IL-38, 10 ng/ml IL-1β, 20 ng/ml IL-38 + 10 ng/ml IL-1β for 6 days. Scale bar: 200 μm. *p < 0.05; **p < 0.01; ****p < 0.0001. Mean ± SEM.

Since we observed that IL-38 induces low concentrations of IL-1β (Figure 6A), we decided to test the response of organoids to different concentrations of this cytokine. As also observed by Katsura and collaborators in lung organoids (31), low concentrations (10 pg/ml) of IL-1β stimulate organoid growth (Figure 6H,I). Notably, higher concentrations do not affect organoid size possibly because of IL-1β-dependent cytotoxicity (Figure 6H,I). The highest IL-1β concentration used (10 ng/ml), on the other hand, probably inducing cell death in the organoids, significantly reduced their size (Figure 6J,K). The protective functions of IL-38 against high levels of IL-1β are confirmed by treating wild type organoids with 20 ng/ml IL-38 and 10 ng/ml IL-1β. Organoids treated with 20 ng/ml IL-38 and 10 ng/ml IL-1β are significantly larger than 10 ng/ml IL-1β-treated organoids and do not appear significantly different to vehicle organoids (Figure 6J,K).

Another confirmation of the protective role of IL-38 was obtained from LPS-treated cultures. LPS exerts detrimental effects on organoids growth (32) (Figure S3A,B). Therefore, we tested three concentrations of LPS (10, 100 and 1000 ng/ml) and we observed that the 10 ng/ml slightly decreased organoid size (p=0.079) compared to untreated organoids, whereas 100 and 1000 ng/ml significantly reduced organoid size compared to untreated controls (Figure S3). To test the protective effects of IL-38, we treated organoids with 100 ng/ml LPS with or without the presence of 20 ng/ml IL-38. Whereas LPS decreases organoid size and IL-38 alone increases size, IL-38 rescues LPS-dependent negative effects (Figure S3C,D).

## Discussion

Several studies highlight the role of interleukins in regulating intestinal crypt functions. For instance, IL-22 protects ISCs from immune-mediated tissue damage and promotes intestinal epithelial regeneration (33, 34). Additionally, IL-22, by activating the STAT3 pathway and inhibiting WNT and Notch signaling, suppresses cell differentiation and intestinal stem cell self-renewal (35). However, a study of human intestinal organoids has recently highlighted the role of IL-22 in the maturation of Paneth cells (36). Similar effects on cell differentiation were observed by Deng and collaborators in their report on IL-10, which leads to inhibition of the WNT pathway and the subsequent depletion of ISCs (37). On the other hand, the chronic exposure of IL-1β induces the expression of stem cell markers like *Bmi1, Lgr5, c-Myc, β-catenin* and *Nanog* in the IEC-18 rat normal epithelial cell line, showing a putative role of this cytokine in stem cell homeostasis (38).

In the present studies, we analyzed the role of IL-38, in order to better define its functions in the gut, including the downstream molecular mechanisms that IL-38 activates to maintain the homeostasis of intestinal tissue. We know that IL-38 reduces intestinal inflammation, and the lack of this cytokine hampers gut recovery after DSS-induced injury (2,7,8). Therefore, we focused on the stem cell compartment in intestinal crypt. We demonstrated that exogenous IL-38 stimulation induces the activation of WNT pathway triggering the secretion of the agonist WNT3a, which results in the increase of organoid size and budding. On the other hand, genetic ablation of IL-38 resulted in the downregulation of several stem cell markers (*Fzd5, Ephb2, Olfm4*). These markers are usually related to stemness and WNT activity (39, 40, 41). In addition, IL-38 deficiency also severely dampened the expression of markers of Paneth cells (*Lyz1*), absorptive cells (*Si*), and goblet cells (*Tff3* and *Chga*). These observations reveal the central role of IL-38 in maintaining the homeostasis of ISCs. In its role in intestinal homeostasis, IL-38 determines the differentiation towards several intestinal cell types, thus explaining the importance of this cytokine in the correct recovery of intestinal injuries.

In order to identify the molecular mechanisms of IL-38 in the intestines, we studied IL-1R9 and IL-1R6, which were identified as putative receptors of IL-38 (5, 10, 12, 13). Intestines taken from IL-38 deficient mice show a significant downregulation of *Fzd5*, *Ephb2, Bmpr1a, Pdgfa, Smoc2, Socs3, Cdk6, Clca4, Olfm4, Phlda* and *Pla2g4a* compared to wild type. Conversely, only *Fzd5, Bmpr1a*, *Phlda* and *Cdca7* are the only stem cell niche markers that appeared significantly downregulated in IL-1R9 deficient mice compared to wild type. *Fzd5* encodes for a Frizzled5 receptor for WNT ligands and crypt proliferation requires it and/or Frizzled8 (42). BMP signaling is essential for driving the correct differentiation of intestinal stem cells and an increasing gradient of BMP from the crypt to the top of the villus determines the proper zonation of intestinal epithelium (43). Among the several receptors of BMP, the one encoded by *Bmpr1a* is important because it regulates the phosphorylation of SMAD1/5/8 in the intestine (26). *Phlda* marks epithelial stem cells (44)*. Cdca7* is another marker of intestinal stem cells (26). The downregulation of these four transcripts in *Il1rapl1^-/y^* intestines reveal a general disorganization of the intestinal tissue in these mice, but not as dramatic as shown in *Il1f10* knockouts. Moreover, we observed that, using the IL-1R9 deficient mouse, the absence of IL-1R9 does not affect IL-38-dependent increase of organoid size, although IL-1R9 deficiency *per se* gives rise growth defects. These findings exclude IL-1R9 as a co-receptor for IL-38, in the intestinal crypt compartment. On the other hand, IL-1R6 silencing not only resulted in a significant downregulation of organoids size and expression of two stem cell markers (*Fzd5* and *Ephb2*), and the lack of IL-1R6 also blocks the responsiveness of organoids to recombinant IL-38 (Figure 5). We conclude that IL-1R6 works as IL-38 receptor in the gut stem cell niche compartment.

Another controversial aspect of IL-38 is its nature as a pro- or anti-pathological cytokine. In most diseased conditions, IL-38 exerts an anti-inflammatory function, yet it can induce detrimental effects (10, 45). As regards colorectal cancer, IL-38 was reported to inhibit the ERK pathway thus suppresses cell migration and proliferation (46). Therefore, given its bipolar nature, IL-38 can be considered as a homeostatic factor that balances the intestine according to its needs. This may also explain why IL-38 works only in a precise concentration range. Upon testing of three different concentrations of IL-38 (2, 20, and 200 ng/ml), only 20 ng/ml induced organoid growth, whereas 2 ng/ml was not sufficient to trigger the IL-38-dependent mechanisms. On the other hand, 200 ng/ml could have saturated the whole receptors necessary for IL-1 signaling, thus inhibiting the effects of IL-1 family members downstream to IL-38. The homeostatic nature of IL-38 was further validated in response to IL-1β stimulation. In fact, in healthy conditions, IL-38 induces low levels of IL-1β (about 20 pg/ml) that helped organoids growth. These findings are in line with the discovery of detectable yet low resting circulating concentrations of IL-1β in healthy patients (47) that are likely necessary to keep the biological systems in a steady state. However, based on our observations in intestinal organoids challenged by high concentrations of IL-1β or LPS, IL-38 exerts a protective signal that restores the normal conditions and acts as an anti-inflammatory cytokine.

All in all, we can conclude that IL-38, by signaling via binding IL-1R6 but not IL-1R9, favors stem cell renewal and differentiation by inducing WNT3a and regulating the expression of crypt-related genes, acting as a growth factor. In this context, IL-38 induces a homeostatic release of IL-1β (at pg/ml level) that supports growth. On the other hand, in inflamed conditions, IL-38 heals injuries by returning the system to the normal state, again exerting pro-homeostatic functions. These observations allow us to conclude that, in the intestine, IL-38 should not be categorized as a pro- or anti-inflammatory cytokine, as in different contexts it can either induce or repress IL-1β to maintain intestinal homeostasis. These findings are remarkably important if we consider the emerging therapeutic potential of IL-38 (14, 15, 16). Our findings revealed that, comparably with what was observed with IL-37, another enigmatic IL-1 family member (48), IL-38 exerts its functions on organoids at a specific concentration. IL-37 and IL-38 are mostly described as anti-inflammatory cytokines (49, 50) and recent studies demonstrated that they both can inhibit trained immunity (51, 52). In conclusion, our results highlight the importance of IL-38 not only as a growth factor, but also as a cytokine that, working only in a specific concentration frame, can be a potent modulator of intestinal inflammation in human therapies.

Furthermore, these data highlight the commercial potential of IL-38 as a supplement of commercially available kits for organoids growth: the addition of IL-38 in these kits would increase the culturing efficiency, accelerating growth and crypt budding.

## Materials and Methods

### Intestinal crypt isolation and organoid culture

Intestinal segments were collected from wild type, *Il1f10* deficient, *Il1rapl2* deficient, and *Nlrp3* deficient mice and were flushed in cold PBS. 2 cm long Intestinal pieces were cut longitudinally and washed vigorously for 20 times to remove mucus and debris. Subsequently, tissues were incubated with Gentle Cell Dissociation Reagent (Stemcell Technologies) in a rocking platform for 20 minutes at room temperature. Cells were removed and intestinal pieces were resuspended in PBS + 0.1% BSA and pipetted up and down 10 times. Supernatant was passed through a 70 μm strainer into a 50 ml tube and centrifuged at 290 x *g* for 5 minutes at 2-8°C. Cell pellets were resuspended in DMEM/F-12 with 15 mM HEPES and centrifuged at 200 x *g* for 5 minutes at 2-8°C. Isolated crypts in a 50:50 mixture of IntestiCult^TM^ Organoid Growth Medium and Matrigel® and 20 μl of this solution was transferred to each well of a preheated 24-well plate. The plate was then incubated at 37°C for 15 minutes and 550 μl of complete IntestiCult^TM^ Organoid Growth Media was added to each well. Media was changed every three days.

### *In vivo* models

Animal protocols were approved by the University of Colorado Animal Care and Use Committee. *Il1f10* deficient mice (GenBank accession number: NM_153077.2; Ensembl: ENSMUSG00000046845) were generated using CRISPR/Cas9 technology and were previously described in de Graaf *et al*. (7); *Il1rapl1* deficient mice (GenBank accession number: NM_001160403.1; Ensembl: ENSMUSG00000052372) were generated by deleting Exon 3 using CRISPR/Cas9 technology (Cyagen Biosciences) and were previously described in The *et al*. (6); the *Nlrp3* knockout mice (B6.129S6-Nlrp3^tm1Bhk^/J) were purchased from The Jackson Laboratories and previously described in Tengesdal *et al.,* (53); the *Il1r1* knockout mice (B6.129S7^tm1lmx^/J) was kindly provided by Shaikh M Atif.

### Protein extraction and western blotting

SI were collected from wild type, *Il1f10* deficient mice. Tissues were were lysed in RIPA buffer (Sigma) supplemented with protease and phosphatase inhibitors (Roche), centrifuged at 13,000 *g* for 30 minutes at 4°C and the supernatants were obtained. Protein concentration was determined in the cleared supernatants using Bio-Rad protein assay (Bio-Rad Laboratories). Electrophoresis was performed on Mini-Protean TGX 4-20% gradient gels (Bio-Rad Laboratories) and blotted onto nitrocellulose 0.1 μm 145 membranes (GE Water & Process Technologies). Membranes were blocked in 5% rehydrated non-fat milk in TBS-Tween 0.5% for 1 hour at room temperature. Primary antibody for LGR5 (R&D Systems) was used in combination with peroxidase-conjugated secondary antibodies. A primary antibody against β-Actin (Santa Cruz Biotechnology) was used to assess protein loading.

### RT-qPCR

Total RNA was extracted from murine SI and from organoid cultures with TRIzol reagent. cDNA synthesis was performed using High-Capacity cDNA Reverse Transcription Kit (Applied Biosystems) according to manufacturer’s protocol. qPCRs were performed in triplicate with SYBR Green Master Mix (Applied Biosystems) by means of QuantStudio 3 Real-Time PCR System (Applied Biosystem). *18s* was used as internal standard in each sample. The sequences of the primers used are listed in Table 1.

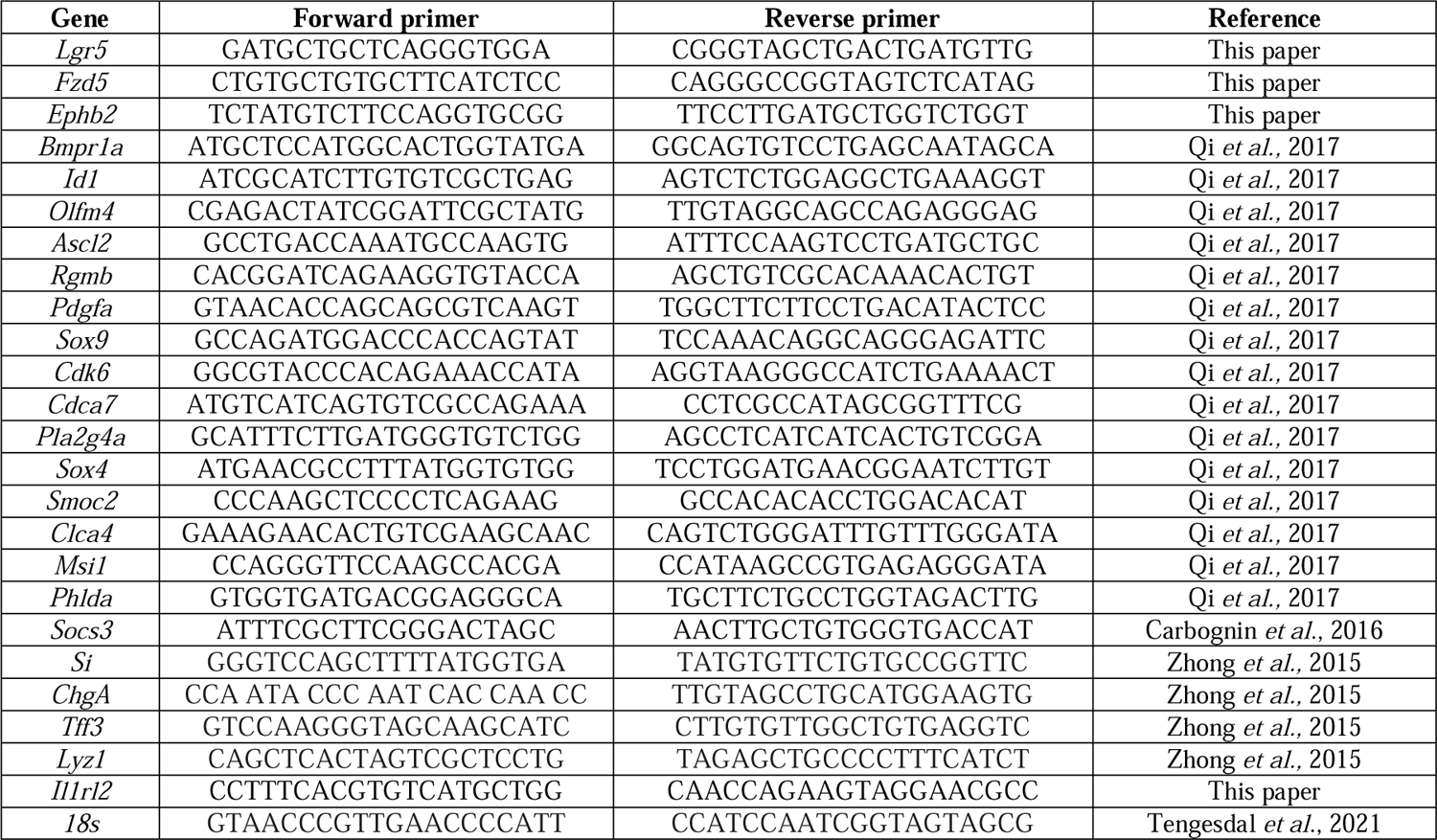

### Imaging

Organoid stacks were taken with Olympus IX81 spinning disk after 6 days of culturing. Organoid size was measured using Fiji (ImageJ) software in pixel^2^, every measurement was normalized with the average of the control group for each experiment.

### Immunofluorescence

Mouse intestinal samples were collected and fixed in PBS with 4% paraformaldehyde (Sigma-Aldrich) in PBS. After dehydration, samples were embedded in paraffin molds for sectioning. Sections (5µm) were obtained and transferred to glass microscope slides. Slides were deparaffinized with xylene and ethanol and permeabilized with 0.1%Tween in TBS solution. After antigen retrieval with citrate buffer, slides were blocked with normal donkey serum and incubated with primary antibodies against LGR5 (rat, R&D Systems) and IL-38 (rabbit, Abcam) dilution overnight at 4°C. Alexa flour 647 anti-rat, Alexa fluor 555 anti-goat and Alexa fluor 488 anti-rabbit at 1:100 (Invitrogen) were used as secondary antibodies for 1h at room temperature. Finally, slides were coverslipped in Mowiol mounting media with DAPI. Images were taken using Olympus FV1000 laser scanning confocal/CARS microscope.

Mouse intestinal organoids were collected, and the pellet wash washed three times with PBS to remove the Matrigel. Then the organoids were fixed in 4% PFA in PBS. After three washes in PBS 0.1%BSA 0.2% Triton X-100 0.1% Tween 20 (Immunofluorescence buffer) to remove the residual PFA, organoids were resuspended in PBS and, subsequently, incubated at 98°C in citrate buffer for 20 minutes for antigen retrieval. After three washes in Immunofluorescence buffer, organoids were permeabilized with a 1% Triton X-100 5% donkey serum PBS for 6 hours. After permeabilization, organoids were incubated with Lysozyme 1 (mouse, Santa Cruz Biotechnology), IL-38 (rabbit, Abcam) at 1:500 dilution overnight at 4°C. The next day, organoids were washed three times with immunofluorescence buffer and incubated with Alexa fluor 555 anti-goat, Alexa fluor 555 anti-mouse and Alexa fluor 488 anti-rabbit at 1:1000 diluted in immunofluorescence buffer supplemented with 10% donkey serum for 2 hours at room temperature on a tilting platform. Secondary antibody solutions were removed with three washes in immunofluorescence buffer and organoids were transferred to a 24 well-plate and organoid Z stacks were taken with Olympus IX81 spinning disk.

### Cytokine Measurements

Cytokine concentrations were measured by specific DuoSet ELISAs according to manufacturer’s instructions (R&D Systems).

### *Il1rl2* silencing

*Il1rl2* siRNAs (MBS8235553) and unspecific siRNA as negative control (MBS8241404) were purchased from MyBioSource and transfected with the siTran 2.0 siRNA transfection reagent following manufacturer’s instructions. For transfection we used 10 nM siRNAs for 24 hours. Silencing efficiency was evaluated with RT-qPCR using the *Il1rl2* primers listed in Table 1.

### Statistical analysis

Significance of differences was evaluated with Student’s *t*-test or using GraphPad Prism (GraphPad Software Inc.). Statistical significance was set at *p* < 0.05.

