## Supplemental data for "IL-38 regulates intestinal stem cell homeostasis by inducing WNT signaling and beneficial IL-1β secretion"

**Supplementary**

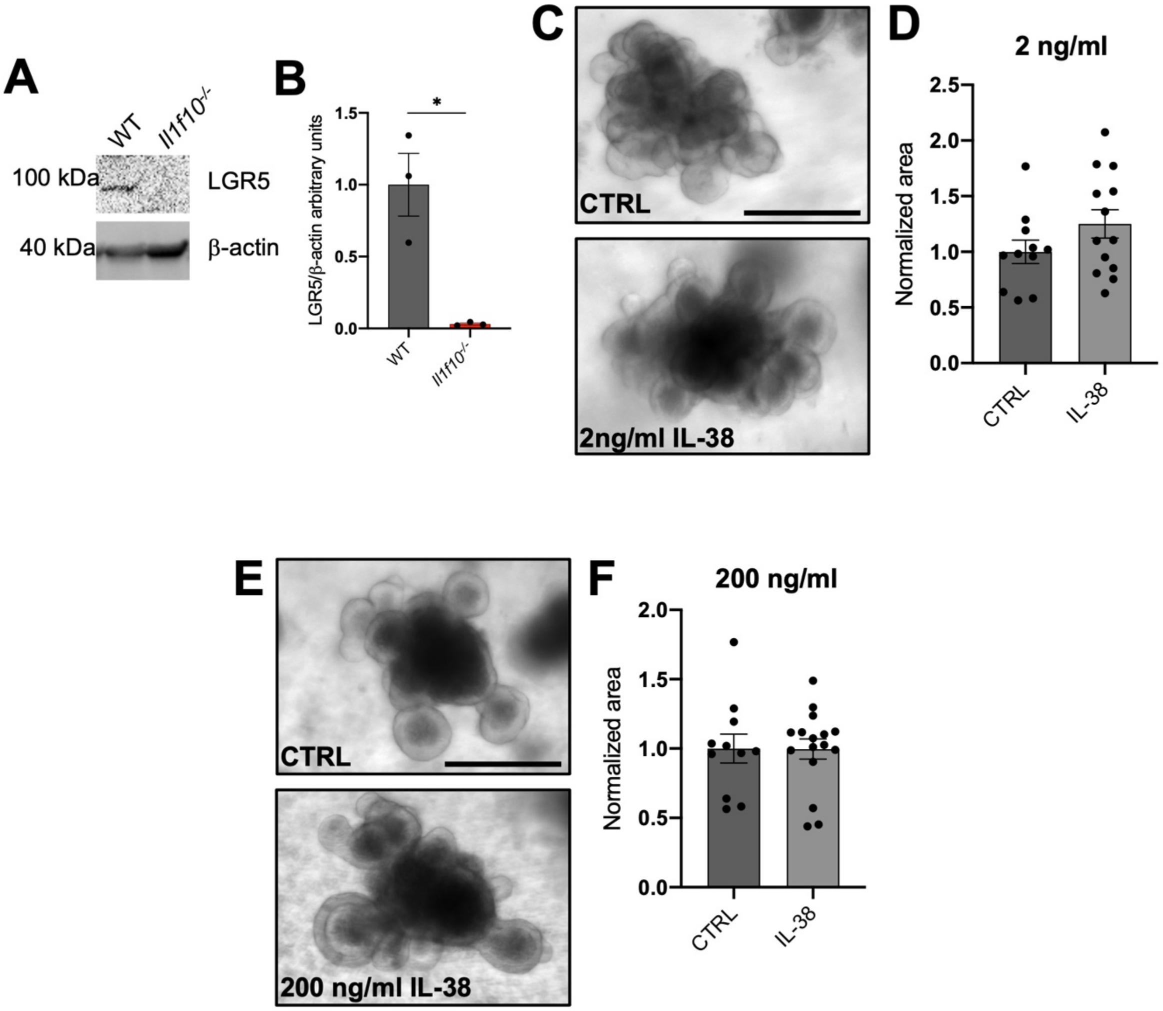


Figure S1: **2 and 200 ng/ml IL-38 do not affect organoid growth.** (**A**) A representative western blot performed on wild type and *Il1f10* knockout intestines. (**B**) Histogram derived from western blot shown in (A) of LGR5 levels in wild type and *Il1f10* knockout intestines. (**C**) Representative pictures of organoids treated with vehicle and 2 ng/ml IL-38. (**D**) Measurements of organoids treated with vehicle and 2 ng/ml IL-38. (**E**) Representative pictures of organoids treated with vehicle and 200 ng/ml IL-38. (**F**) Measurements of organoids treated with vehicle and 200 ng/ml IL-38. * $p < 0.05$. Mean ± SEM.

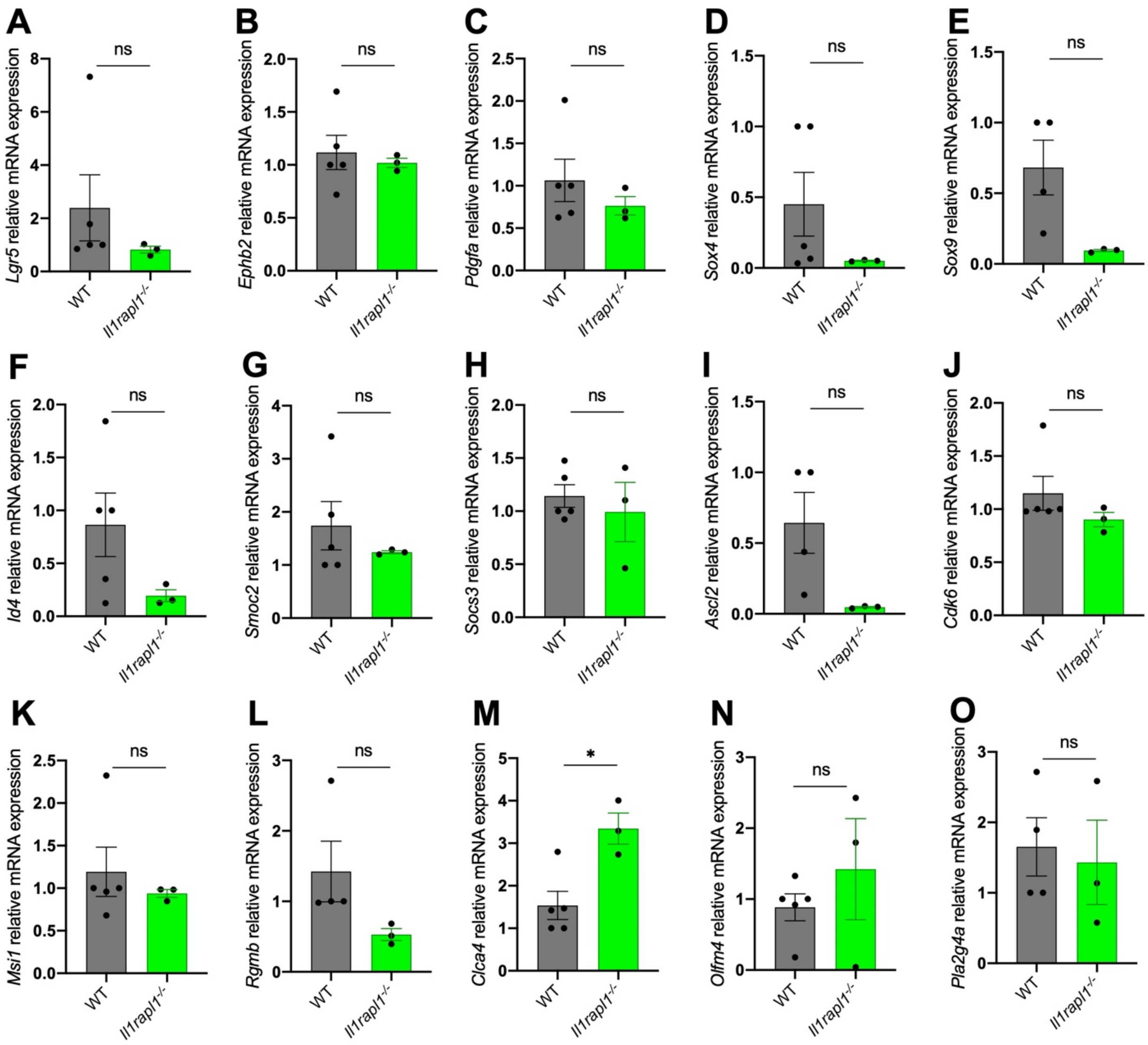


Figure S2: **Crypt markers in IL-1R9 KO intestines**. (**A-O**) Gene expression analysis of *Lgr5* (**A**), *Ephb2* (**B**), *Pdgfa* (**C**), *Sox4* (**D**), *Sox9* (**E**), *Id4* (**F**), *Smoc2* (**G**), *Socs3* (**H**), *Ascl2* (**I**), *Cdk6* (**J**), *Msi1* (**K**), *Rgmb* (**L**), *Clca4* (**M**), *Olfm4* (**N**) and *Pla2g4a* (**O**) in intestines from wild type and *Il1rapl1* deficient mice. * $p < 0.05$, ns = not significant. Mean ± SEM.

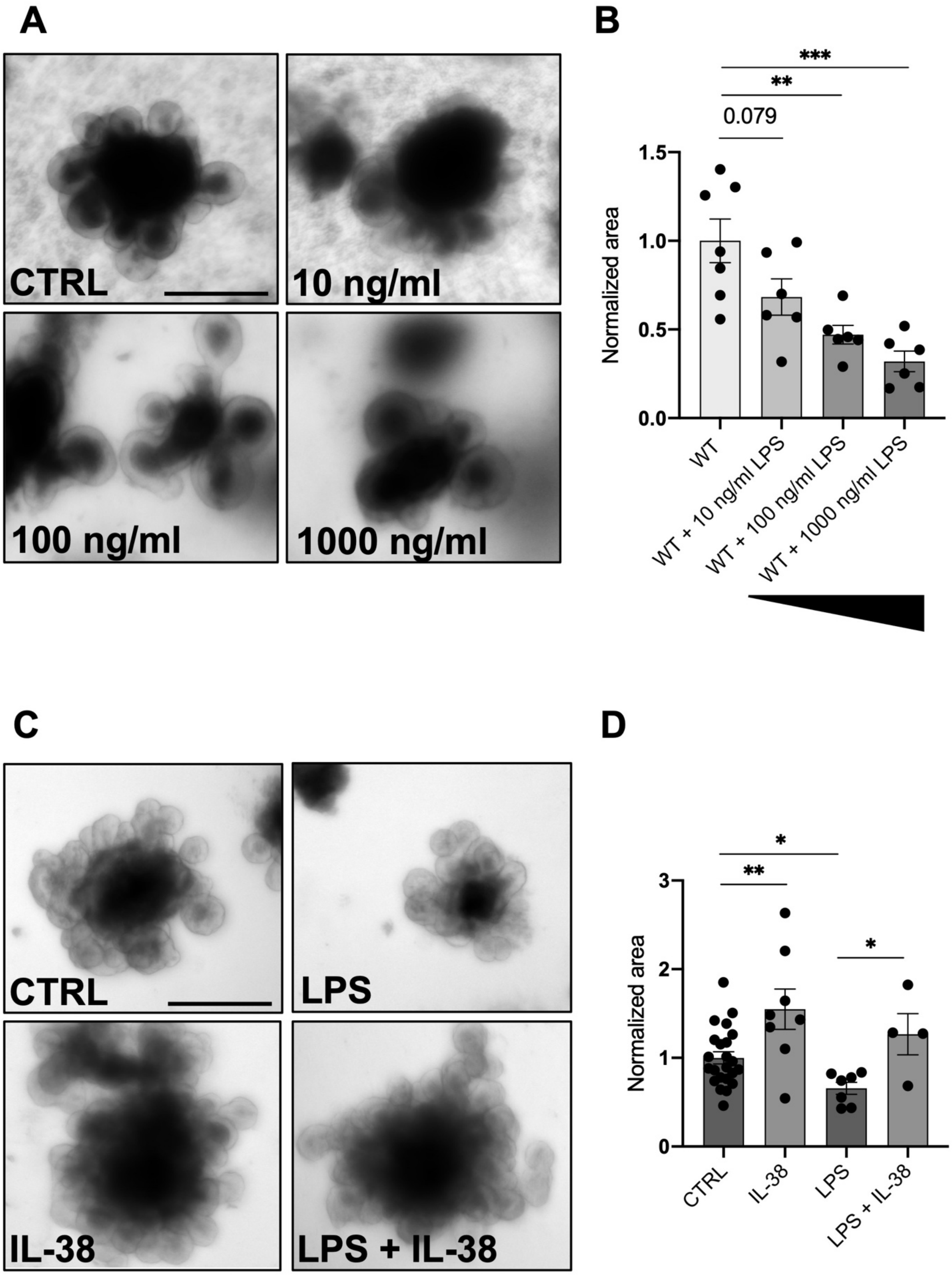


Figure S3: **LPS has negative effects on organoid growth.** (**A**) Representative pictures of wild type organoids treated with vehicle, 10 ng/ml, 100 ng/ml and 1000 ng/ml LPS for 6 days. (**B**)

Measurements of wild type organoids treated with 10 ng/ml, 100 ng/ml and 1000 ng/ml LPS for 6 days. (**C**) Representative pictures of wild type organoids treated with vehicle, 20 ng/ml IL-38, 100 ng/ml LPS, 20 ng/ml IL-38 + 100 ng/ml LPS for 6 days. (**D**) Measurements of wild type organoids treated with vehicle, 20 ng/ml IL-38, 100 ng/ml LPS, 20 ng/ml IL-38 + 100 ng/ml LPS for 6 days. *$p < 0.05$; **$p < 0.01$; ***$p < 0.001$. Mean ± SEM.
